# Proteomics of phosphorylation and protein dynamics during fertilization and meiotic exit in the *Xenopus* egg

**DOI:** 10.1101/145086

**Authors:** Marc Presler, Elizabeth Van Itallie, Allon M. Klein, Ryan Kunz, Margaret L. Coughlin, Leonid Peshkin, Steven P. Gygi, Martin Wühr, Marc W. Kirschner

## Abstract

Fertilization triggers release from meiotic arrest and initiates events that prepare for the ensuing developmental program. Protein degradation and phosphorylation are known to regulate protein activity during this process. However, the full extent of protein loss and phospho-regulation is still unknown. We examined absolute protein and phospho-site dynamics after fertilization by mass spectrometry-based proteomics. To do this, we developed a new approach for calculating the stoichiometry of phospho-sites from multiplexed proteomics that is compatible with dynamic, stable and multi-site phosphorylation. Overall, the data suggest that degradation is limited to a few low abundance proteins. However, this degradation promotes extensive dephosphorylation that occurs over a wide range of abundances during meiotic exit. We also show that eggs release a large amount of protein into the medium just after fertilization, most likely related to the blocks to polyspermy. Concomitantly, there is a substantial increase in phosphorylation likely tied to calcium activated kinases. We identify putative degradation targets as well as new components of the block to polyspermy. The analytical approaches demonstrated here are broadly applicable to studies of dynamic biological systems.

**Significance Statement:** Protein phosphorylation and degradation drive critical events in early embryogenesis and the cell cycle; however, comprehensive and accurate analysis of these changes is currently difficult. Using a mass spectrometric approach, we present a quantitative view of the protein and posttranslational economy of the fertilization response in the frog egg. Protein degradation affects a small but very important class of proteins, while regulatory phosphorylation and protein release occur on a far larger scale. We have developed new, broadly applicable analytical methods for phosphorylation that provide absolute quantification with confidence intervals for improved interpretability of post-translational modification analysis.

## Introduction

For decades, the highly synchronous events of fertilization have provided a useful system for the study of many aspects of cellular regulation, especially protein degradation and phosphorylation. The destruction of Cyclin-B and other proteins is catalyzed by the anaphase promoting complex (APC/C), which promotes M-phase exit (1). The two activators of the APC/C are Cdc20 and Cdh1. In the egg, the cell cycle typically involves only Cdc20 (2). While the list of known Cdh1 substrates continues to grow (3), the Cdc20 target list remains small (4–6). It is of considerable interest to characterize the minimal set of Cdc20 substrates that powers the early cell cycle. Kinase activity is equally important to this regulation. Cyclin-B degradation promotes mitotic exit by inhibiting the activity of Cyclin-dependent Kinase 1 (Cdk1). There is a bulk loss of phosphorylation following egg activation (7). The identities of the vast majority of these phosphoproteins remain undiscovered. However, there is a strong expectation that this regulation reflects the reversal of Cdk1 phosphorylation. Fertilization employs additional regulation not common to other cell cycles. There are timed waves of phosphorylation that correspond to the release of cortical granules just after fertilization as part of the slow block to polyspermy. This release is calcium sensitive, and may reflect increases in the activity of Protein Kinase C (8, 9) and CaMKII (10). An account of the secreted proteins, their function, and their upstream signaling components is limited. To investigate these unknown aspects of degradation, release, and modification of proteins at fertilization comprehensively, we employed mass spectrometry (MS) in *Xenopus laevis* eggs. Recent advances in multiplexed proteomics allow quantitative comparisons of multiple conditions using Tandem Mass Tags (TMT) (11–13). Our recent work demonstrates the power of proteomics in *Xenopus* for studies of early development (14–16). Though fertilization has been studied before with MS (17–20), the studies were either qualitative or did not measure phosphorylation. With an enrichment step (21–23), it is feasible to measure relative changes in a large number of phosphoproteins. However, such studies are of limited utility without measuring site stoichiometry. Relying on fold change alone for phosphorylation studies will not distinguish between ten-fold relative changes from 1% occupancy to 10% versus 10% to 100%. Several approaches are available (24–26), but none of these are able to determine occupancies of peptides measured with multiplexed MS or that have multiple phosphorylated residues. This paper introduces new biological findings about fertilization and cell cycle progression, but it also introduces new methods for measuring absolute stoichiometry of phospho-sites, widely applicable to MS protein modification studies.

## Results

### Quantitative proteomics of egg activation and meiotic exit

The large (1.2 mm) *Xenopus laevis* egg offers superb synchrony and sufficient material (~30μg of non-yolk protein per egg (27)) for proteomic and phosphoproteomic studies. To capture the progression of the rapid fertilization response (Fig. 1A), metaphase-arrested eggs were activated with electro-shock and snap-frozen every two minutes in sets of 30 (Fig. 1B). We chose electrical activation over sperm addition to maximize time resolution by avoiding the asynchrony of sperm penetration through the jelly coat (28). The early morphological and molecular events to our knowledge are equivalent between fertilization and parthenogenetic activation (29, 30), hence we use the terms interchangeably here. More than 99% of eggs activate with a standard deviation of <15 sec using this approach (SI Appendix, Fig. S1 and Movies 1-4). Time points were analyzed by MultiNotch MS3 (31). Phosphopeptides were enriched on a titanium dioxide column; the flow-through was used for protein-level MS. We multiplexed the TMT-labeled time points before enrichment to eliminate errors that arise from variation between columns. Protein-level MS was performed with four biological replicates, and phospho-level MS was performed in biological triplicate. We quantified ~8,700 proteins (80% detected in two or more replicates) and ~3,500 phospho-sites (40% in two or more replicates) on 1,700 proteins (Fig. 1C). Less than 1% of detected proteins showed abundance changes. 40 proteins decrease in abundance over time, while 8 unexpectedly show an apparent increase (Fig. 1D) (discussed below). Protein phospho-sites are notably more dynamic, with ~15% of modified residues changing (Fig. 1E). There is clear evidence for parallel dephosphorylation and phosphorylation. The phospho-site changes overwhelmingly occur on stable proteins, thus they reflect actual changes in phosphorylation level rather than protein level.

**Figure 1:**
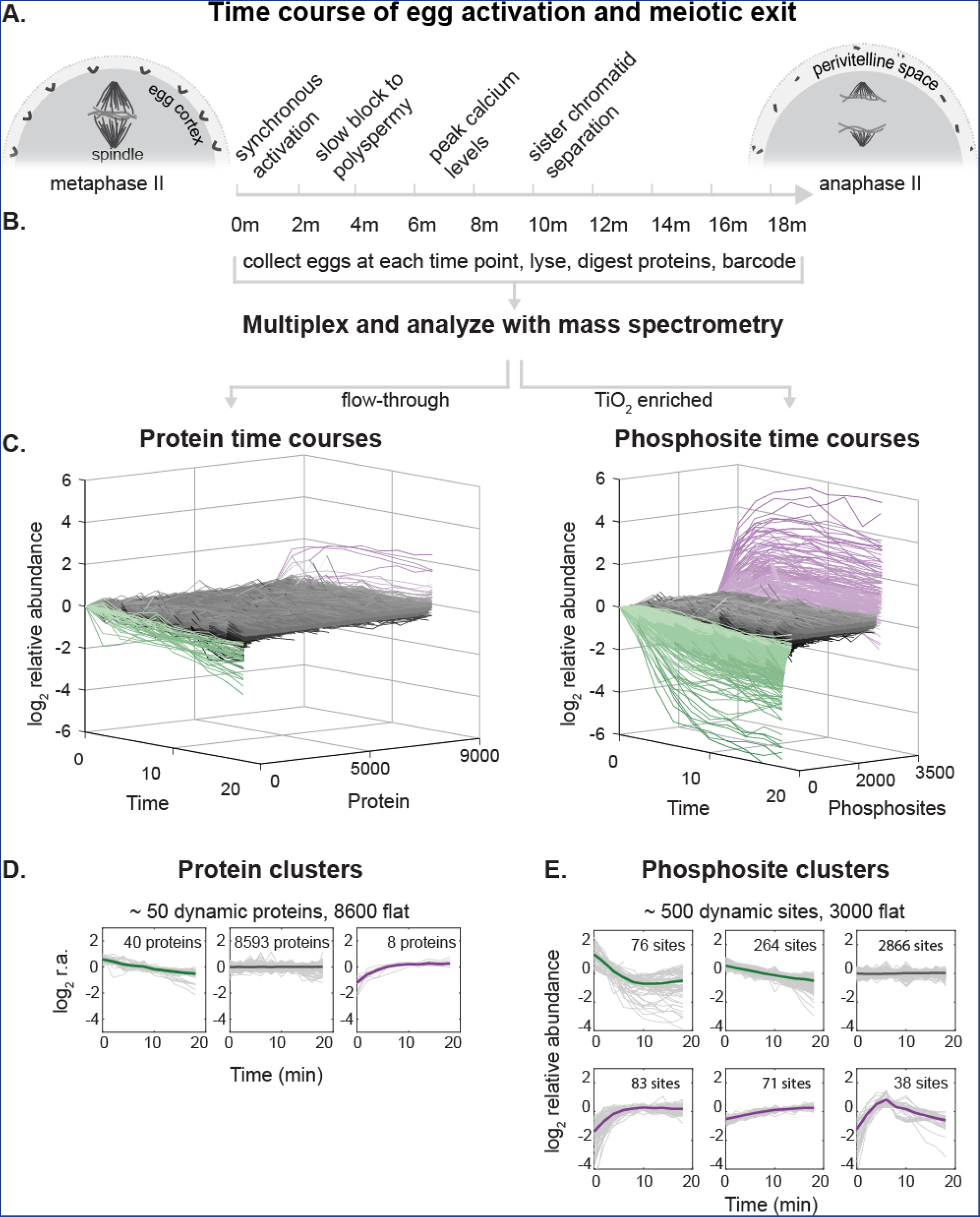
Experimental overview. A) Fertilization and egg activation release metaphase-arrested oocytes into anaphase and initiate the slow block to polyspermy, sister chromatid separation, remodeling of the specialized egg extracellular matrix and inflation of the perivitelline space, among other processes. We mimic fertilization and trigger egg activation via electric shock to maximize synchrony. B) After activation, eggs were collected every two minutes, lysed, and digested with proteases. Samples were barcoded using Tandem Mass Tags (TMT) and multiplexed. Protein and phospho-site dynamics were measured with mass spectrometry-based proteomics. Phosphorylated peptides were enriched using TiO_2_, and the column flow-through peptides were used for protein analysis. C) Modified waterfall plots displaying the trend of every protein and phospho-site in the dataset, normalized to the first time point and ordered first by clusters (see below) and then in ascending order within each cluster. D, E) K-means clustering to summarize dynamic classes of the protein (D) and phosphorylation time series (E) (see SI Appendix, Methods). Bold line represents the centroid of the cluster while the gray lines are individual time courses (normalized to the time course means).

### Protein loss following fertilization

We sought to determine whether additional proteins beyond the known cell cycle targets are degraded. We identify 29 unique proteins by gene symbols that significantly decrease in abundance (Table S1, Table S2). To establish statistical significance, we calculated a randomization-based False Discovery Rate (FDR) (SI Appendix, Fig. S2) and set a cutoff of 1%. Specifically, we find known APC/C^Cdc20^ substrates, including Cyclin-B1, Cyclin-B2, Geminin (32), Securin (33, 34), and the *β*-TRCP substrate EMI2 (35) decrease significantly (Fig. 2A). Previously, we estimated the absolute abundances of ~11K proteins in the frog egg (14), which allowed us to estimate absolute rates of degradation for these proteins. We see a small delay of ~2- 4 minutes between fertilization and initiation of degradation of the APC/C substrates. EMI2, whose destruction is required for APC/C activity, declines without delay. It is reduced by 30% 4 minutes after activation, when Cyclin-B1/2 and Securin begin to decline. EMI2 is stoichiometric with the least abundant components of the APC/C (14). Therefore, a ~30% decline of EMI2 leads to a 30% increase in the maximal activity of APC/C. This activity must therefore be sufficient to initiate degradation. All observed putative APC/C^Cdh1^ substrates, such as CDC20 (36–38), PLK1 (5), and Aurora A/B (39, 40) (SI Appendix, Fig. S3), are stable. The total loss of the known APC/C^Cdc20^ substrates is ~3,000 pg (or 250 nM), and they are degraded at a rate of ~150 pg/min. From single molecule measurements of the proteasome (41), we estimate the reactions responsible for protein degradation are approximately diffusion-limited for the reaction (see Methods). Since it is possible to competitively delay substrate degradation by adding peptide fragments of Cyclin B (32, 42–44), the reaction in the egg may be near saturation. The overall estimated rates may be affected by the cell cycle metachrony (i.e., spatial heterogeneity) of the large *Xenopus* eggs. The cell cycle state should equilibrate between the top and bottom of the egg by ~20 minutes given the speed of the traveling “waves” (45–47) and other previous measurements (48). Therefore, the end points we report are likely to be accurate. The half times may be overestimated if there is substantial delay of degradation in the vegetal pole, though the exclusion of the cytoplasm by yolk may partially compensate for this phenomenon.

**Figure 2:**
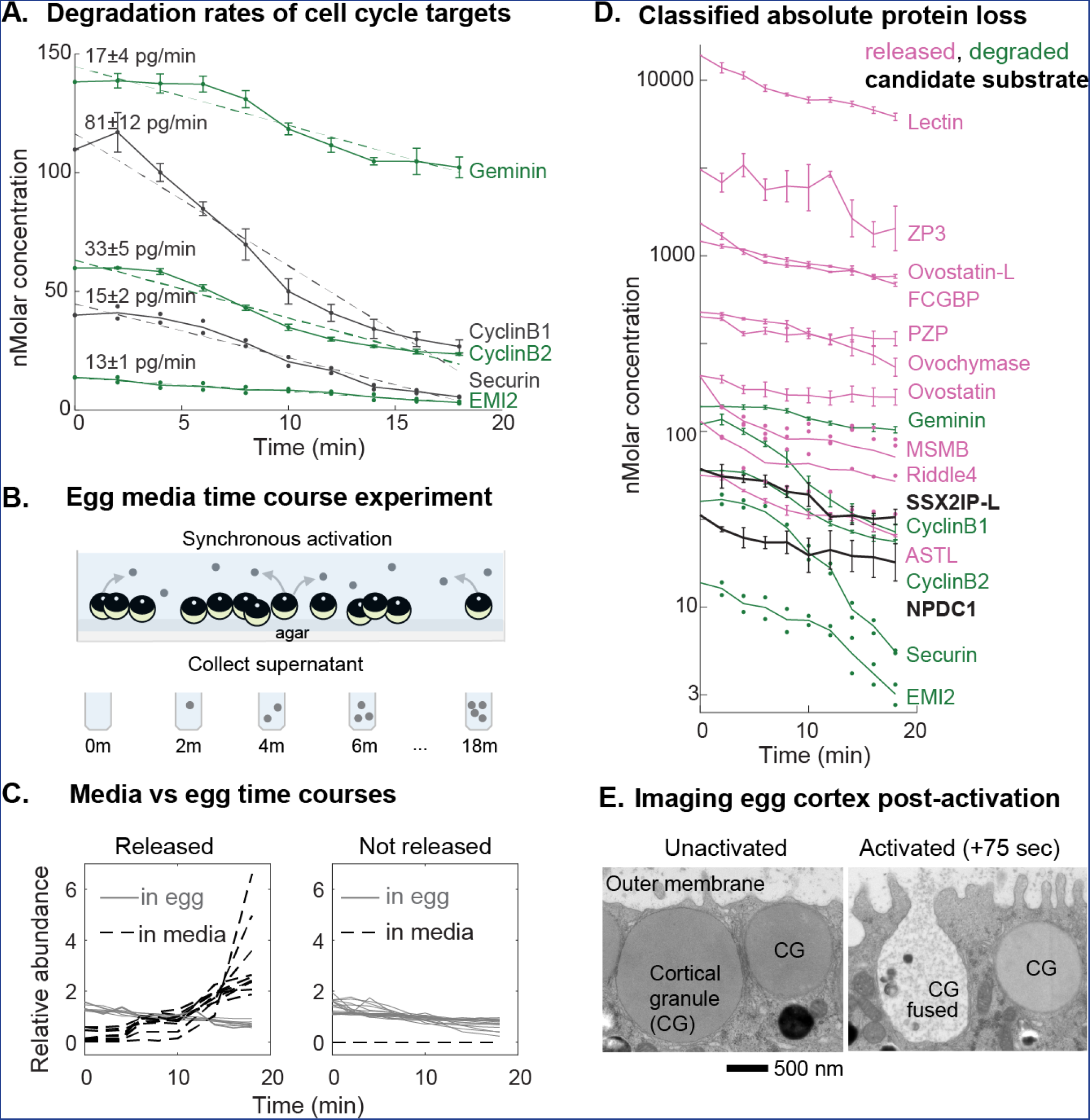
Proteins loss occurs by degradation and release. Experiments and analysis performed to classify the mechanism of protein loss following egg activation (see text). A) Time series of the 5 proteins identified as significantly decreasing which are known cell cycle degradation targets, plotted by their absolute changes. Error bars reflect standard error (SEM) for proteins detected in at least three biological replicates. For proteins detected in fewer than three replicates, all data available is shown as points. Dashed lines represent a linear fit to the approximate zero-order kinetics (labeled with the slope and 95% confidence interval). B) Experimental design to test for proteins released by eggs upon fertilization as an explanation for protein loss. Released proteins will increase in the media fraction over time. C) Time series for proteins detected in both the egg and supernatant or the egg alone. These data were used to classify whether proteins were lost by release rather than degradation (by direct evidence or annotation, see text). This class comprised all but two of the significantly decreasing proteins (see 2D) besides the known targets shown in 2A. D) Time series of protein loss plotted by their absolute abundance (log10 transformed) and classified by the mechanism of loss. Error is visualized as in 2A. SSX2IP-L and NPDC1 homologs (black) are putative new degradation targets (see text for full criteria). E) Electron microscopy images of the cortical granules (CG) on the egg cortex. After egg activation, cortical granules fuse with the outer membrane and expel their contents.

### Classifying candidate novel degradation targets

Annotation of the decreasing proteins suggested that some candidate proteins were released from the cell rather than degraded (Table S2). To test this hypothesis, we harvested the egg media in time series after egg activation (Fig. 2B). Proteins that appeared in the media and disappeared from the cell were classified as released (Fig. 2C and SI Appendix, Fig. S4). Release from the egg is the major source of protein loss rather than degradation (Fig. 2D). We found only six proteins that decreased abundance with no evidence for release (see Table S3 and Methods section for full classifications and rationale). To determine whether these proteins are degraded through the ubiquitin-pathway, we performed ubiquitin pull-downs with a di-glycine antibody (49) and analyzed the samples by MS. Of the six proteins that were not previously known degradation targets, two were ubiquitinated (Fig. 2D, black). A *Xenopus* homolog of the gene SSX2IP (sequence provided in Table S1, distinguished as SSX2IP-L) is the most promising candidate for a novel APC/C substrate (Fig. 2D). SSX2IP is a centrosomal protein (50) with roles in the primary cilium (51). It is present at similar abundance to Cyclin B (60nM) and contains a confident D-box prediction (4). Ciliary defects occur when Cdc20 is depleted (52). SSX2IP may provide a mechanism for linking Cdc20 to the ciliary phenotypes. The other candidate is NPDC1, a neuronal protein with some association with cell cycle components (53) containing a weak D-box motif. However, it is localized to synaptic vesicles in neurons, which may suggest association with exocytosis machinery rather than degradation (though we do not detect it in the media). At the level of sensitivity in this study, the number of putative APC/C^Cdc20^ substrates is indeed very small. All of them are below 100nM in the egg. To estimate the sensitivity of our study, we compared the proteins identified using TMT-MS to those identified by the more sensitive (but semi-quantitative) label-free approach in the frog egg (14) (SI Appendix, Fig. S5). At 50nM, we detect 95% of the proteins with TMT-MS that we do with label-free, and still detect 50% at 10nM. While we cannot rule out additional substrates, the data does suggest that any we did not detect are at low concentrations.

### Proteins released from the egg

By contrast to the relatively small mass and small number of proteins degraded after fertilization, there is a substantial reduction in cytosolic protein due to release into the medium. We detect the expected cortical granule components like an Interlectin2 homolog, as well as the proteases Ovostacin (54) and Ovochymase (55) whose function is thought to be inhibition of further sperm binding by cleaving the extracellular Zona Pellucida (ZP) gene family (56). We detect ZP2, 3 and 4 homologs (57) as released, which are likely to be cleaved peptide fragments that have diffused into the media (SI Appendix, Fig. S4). Therefore, at least two mechanisms are responsible for the release of protein from the interior of the egg and from its surface: exocytosis of cortical granules and proteolysis of the specialized extracellular matrix. We would not detect change for proteins that are trapped in the perivitelline space. We detect several proteins previously unknown to be released from the egg at fertilization, including several protease inhibitors. One examples is Ovostatin (AM2 homolog) (58), which was previously shown to have anti-trypsin activity in the sea urchin egg (59). There are additionally a small number of other annotated protease inhibitors that decrease in the egg but are not detected in the medium. We surmise that these are released as well, rather than degraded (Table S3). One example is Riddle4 (60) with known functions reported later in development (61). While the secretion of proteases is a well-recognized mechanism, the release of endogenous protease inhibitors in response to fertilization was not known. Another previously unknown released protein is a FCGBP homolog (Fig. 2D), which is similar to Zonadhesin (62, 63) that binds to the ZP proteins. It is likely extracellular and liberated by proteolysis. To compare the measured release of protein with the major morphological events of fertilization, the cell cortex of activated eggs was imaged by electron microscopy (EM) every 15 seconds (Fig. 2E). The release of cortical granules shows a 60 second delay, but between 75 and 90 seconds ~40% of the vesicles fuse with the outer membrane. By 10 min, the majority (~60%) of the vesicles have fused (SI Appendix, Fig. S6). This is consistent with the generally faster loss rates of released proteins (Fig. 2D) and with previous work (9).

### Observed protein increase is a phosphorylation artifact

A cluster of 8 proteins appear to accumulate post-fertilization (Fig. 1D), 7 of which show a significant increase in abundance after the FDR analysis (Table S4). Many of these proteins are typically stable and not expected to be synthesized during anaphase (e.g., NUP35, a nucleoporin (64)), which led us to search for an alternative explanation. Since protein trends are determined from the sum of all unmodified peptides, we examined the individual peptide trends to determine whether they all showed the same unexpected increase. We found that these proteins did in fact have stable as well as increasing peptides (e.g., SI Appendix, Fig. S7). This discrepancy can occur from the loss of a modification on that particular peptide. The relative abundance of the unmodified peptide must increase if the modified form decreases, due to conservation. Indeed, for three of these proteins, we can directly show the loss of phosphorylation that causes the increase in the unmodified peptide (SI Appendix, Fig. S7). Overall, we have evidence that 6 of the 7 apparently increasing proteins have phosphopeptides showing dramatic decreases in phosphorylation. Therefore, we conclude that the increasing protein trends are most likely caused by dephosphorylation rather than rapid protein synthesis. We controlled for phosphorylation artifacts in protein loss as well, and found evidence that this impacted only two proteins (Table S3). We found that analyzing the data with metrics that reduce the effect of outliers (e.g., median) mitigates but does not eliminate specious trends caused by dynamic modifications. This is because multiple peptides are often affected by modifications (e.g., SI Appendix, Fig. S7C), as is the case for the majority of the proteins discussed here.

### A method for evaluating phospho-site occupancy

The phospho-MS data (Fig. 1E) show the relative abundance changes of ~500 dynamic and ~3,000 stable phosphorylation sites. While relative trends of phospho-forms have utility, they are often difficult to interpret. For example, a 2-fold increase in the occupancy of a residue from 1% to 2% appears identical to the same fold increase from 50% to 100%. These two cases can have very different functional implications, thus we sought to calculate the phospho-site stoichiometry changes in our data (Fig. 3 and Fig. 4). It is not possible to directly determine stoichiometry from simple ratios of MS measurements without expensive spike-in standards (65). This is because the unmodified and modified forms of a peptide ionize at different efficiencies and hence the inter-form ratio is distorted (Fig. 4A) without standards. It is nonetheless possible to infer the absolute stoichiometry of a site by invoking the principle of mass conservation, which states that the total abundance of each peptide equals the sum of all modified and unmodified forms of the peptide (Fig. 3, Eq.1-3). This principle results in one constraint – a simple linear equation – for each pair of conditions in the experiment. If protein levels change, the equations must be scaled accordingly. By solving these conservation equations, we can infer the occupancy ratio from the measured relative intra-form ratios of change between conditions (Fig. 3, Eq.4). Approaches invoking mass conservation were previously reported (24, 66) for the simple case of two phospho-forms and two conditions, giving two linear equations to be solved for the stoichiometries of each form. Here, we extend the solution and broaden its utility: (i) to include cases where the number of conditions is larger than the number of observed phospho-forms; (ii) to establish a statistical measure of confidence in the results; and (iii) to cases with multiple phospho-forms, which were not calculable by previous approaches.

**Figure 3:**
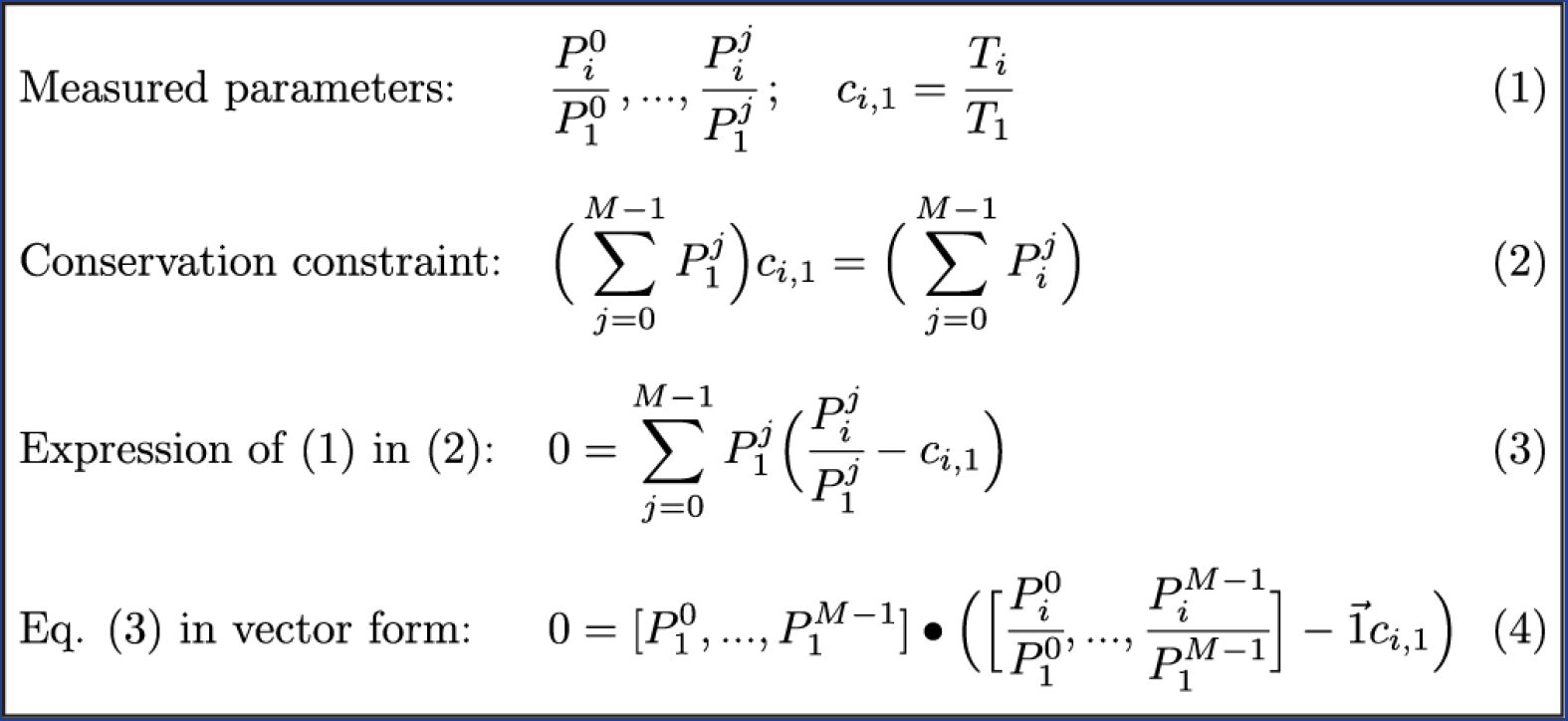
The geometric relationship between the measured peptide form ratios and their unknown absolute values. Let 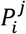 equal the absolute abundance of peptide form *j* in condition *i*, where *j* = 0,1,…,M-1, 0 = unmodified and M is the number of forms, and where *i* =1,…,N, where N is the number of measurements. 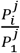 is the MS signal of form *j* at condition *i* normalized by the reference condition *i* = 1. *c_i_*_,1_ is the ratio of the parent protein T between measurement (i) and measurement 1. (Eq.1). The sum of all peptide forms is conserved or scaled by *c_i_*,_1_ 1 (Eq. 2). Rewriting Eq. 2 in terms of the measured parameters (1) yields Eq. 3, which is written in vector form as Eq. 4. Eq. 4 shows that the vector of absolute values is orthogonal to the *M*-1 dimensional subspace containing the measured ratios. When *M* = 2, this subspace is a line (SI Appendix, Fig. S8). For *M* > 2, this subspace is an *M*-1 dimensional plane. For over-determined systems (N > M), this subspace can be estimated with regression (Fig. 4, SI Appendix Methods).

**Figure 4:**
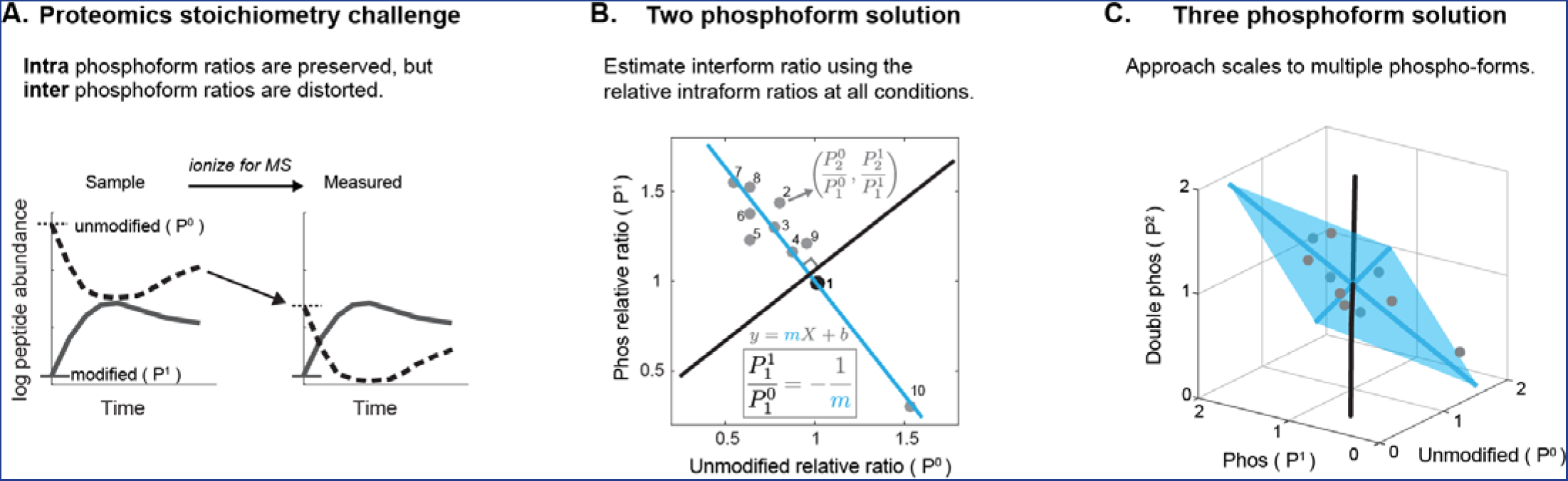
Calculation of phospho-stoichiometry from multiplexed phospho-dynamics measurements. Demonstrating a graphical approach to estimating phospho-site occupancy of multiplexed data. A) Phospho-site occupancy cannot be directly calculated from the raw signal from the mass spectrometer because the inter-form ratios are distorted due to the differential ionization efficiencies of the peptide forms. This is depicted here as the unmodified form (P^0^) ionizing less efficiently than the modified form (P^1^). However, the intra-form ratios are preserved (i.e., only the starting point is shifted). B-C) Estimating the solution for the over-determined system from multiplexed-MS data. B) For each condition, the measured intra-form ratios of change of the unmodified (P^0^) and single phosphorylated (P^1^) forms (see Fig. 3) define coordinates in a 2D plane. The relative ratio is defined by the reference point, which in this case is point 1 (black point). The solution to the over-determined system can be estimated by regression. The unknown inter-form P^1^/P^0^ ratio is the negative inverse of the fit line slope (i.e., is the orthogonal slope. See Fig. 3 (Eq. 4), SI Appendix (Fig. S8), and SI Appendix Methods). C) For more than two P forms, the known ratios define a higher dimensional plane (e.g., a plane is fit in 3D space from the ratios of the P^0^, P^1^, and double form P^2^). The plane is visualized as the blue shaded area spanned by two vectors. The solution (the black vector) is orthogonal to the plane, as in Fig. 4B. These vectors are calculated by principal component analysis (see text). Data points are plotted in grey, blue-shaded points are behind the plane.

Briefly, the method takes advantage of the fact that when the number of conditions exceeds the number of possible phospho-forms, the system (Fig. 3, Eq. 4) is over-determined (SI Appendix, Fig. S8) and the problem becomes one of multivariate linear regression. Thus, an estimated solution to the system can be determined through slope-fitting, as visualized in Fig. 4B and SI Appendix, Fig. S9. This extends to fitting a hyperplane in higher dimensional space for more than two phospho-forms (Fig. 4C). The ability to incorporate many measurements under different conditions enables a more accurate calculation of occupancy than one measurement alone. The “over-determination” of the system is key to reporting confidence intervals. We are able to estimate error in the fit coefficients using bootstrapping (67) (SI Appendix, Methods and Fig. S10). Solving for multi-site occupancy is not possible if the number of conditions is less than the number of phospho-forms, as the system is underdetermined. Therefore, our ability to calculate multisite stoichiometries is enabled by measuring multiple conditions by multiplex-MS.

### Applying occupancy calculation

We have applied this method to the phosphorylation dynamics in the period following egg activation for sites where we detect the unmodified and modified forms. We did not scale the conservation equations in this case since the protein levels are overwhelming stable (Fig. 1C,D). Fig. 5A shows the relative dynamics of phospho-sites on the kinases NEK3 and PAK2, as well as transcription factors/nucleotide-binding proteins SOX3 and YBX2. The amount of relative change for each set is nearly identical. However, YBX2 and NEK3 sites change phosphorylation for only a small fraction of their residues, whereas the residues on SOX3 and PAK2 change substantially. These are selected examples of many important regulatory proteins with discrepancies between relative and absolute changes (SI Appendix, Fig. S11). A proof-of-principle multi-site phosphorylation stoichiometry calculation is shown for a CaMKII-γ peptide (Fig. 5B) with an unmodified (P^0^), single (P^1^), and double phosphorylated (P^2^) form. We estimate the occupancy of the P^0^ form with high confidence; the confidence intervals for the P^1^ and P^2^ forms are wide but non-overlapping, so we are able to conclude that the P^1^ form is more occupied than the P^2^. This highlights the utility of confidence intervals, especially as error increases in higher dimensional spaces. A general consideration when applying this method is that it is most effective under conditions where the relative changes are substantial. For stable sites, the minimization is unreliable and gives wide confidence intervals. To extend the use of this approach for stable sites, we repeated the fertilization time course and artificially induced dynamics with a nonspecific phosphatase treatment of two conditions (25) (SI Appendix, Fig. S12 and Methods). We used a heat labile phosphatase, which allowed us to inactivate the phosphatase activity for subsequent experimental steps. Three examples of calculations of stable occupancies using this method are shown in Fig. 5C. They also demonstrate the range of confidence intervals seen within the data (shown in SI Appendix, Fig. S13B). For corroboration, we used the matching phosphatase treated and untreated time points (25) to calculate occupancy using the previously reported two-condition analytical method (24). We compared these results to the occupancies derived from applying our regression-based approach to the independently measured endogenous dynamics. The results are generally consistent (Pearson coefficient = 0.8).

**Figure 5:**
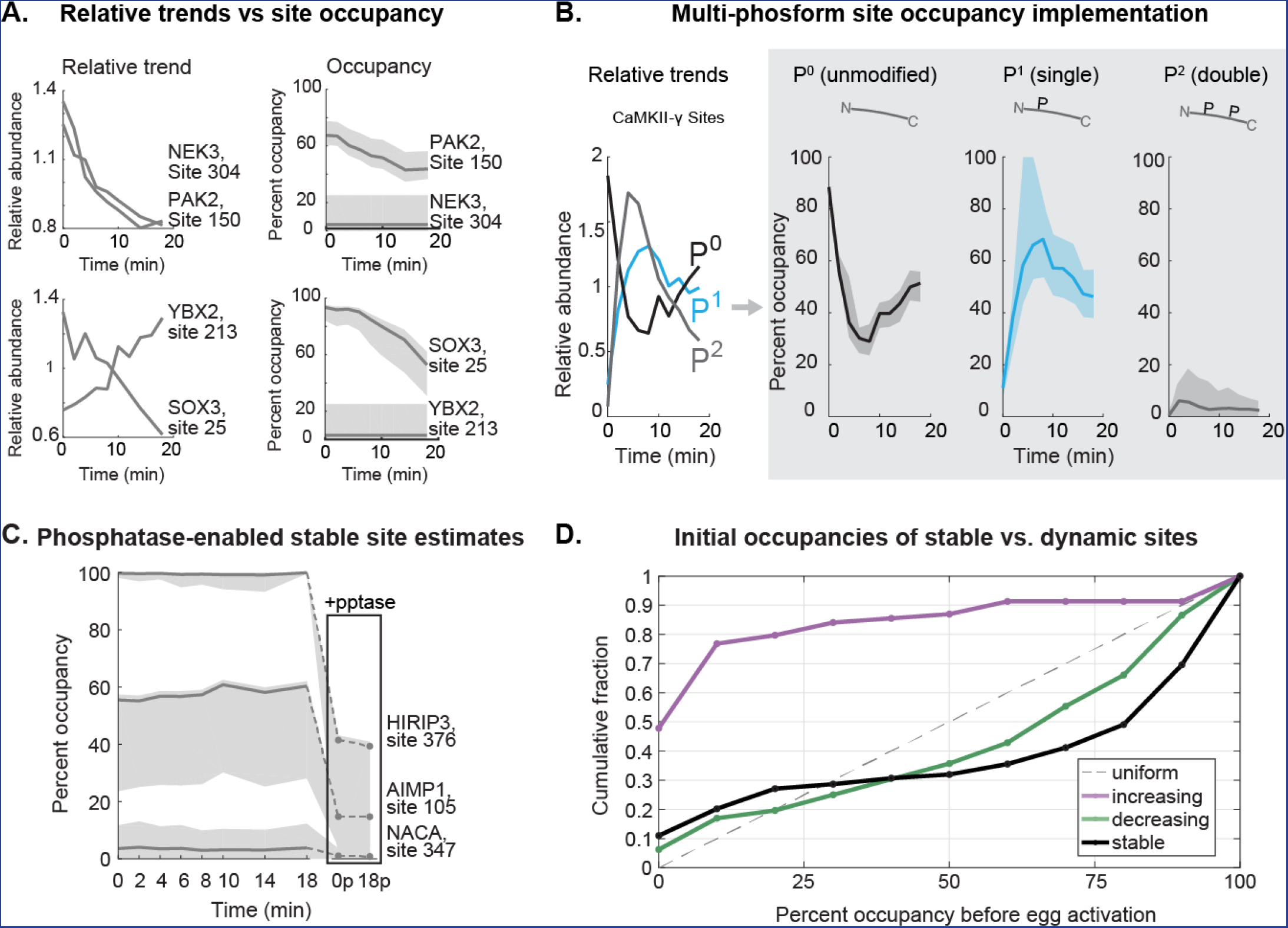
Occupancy of phosphorylation sites in time series post-activation. A) Time series of two kinases (NEK3, PAK2) and two transcription factors/nucleotide binding proteins (SOX3, YBX2) demonstrating that similar amounts of relative changes can give very different occupancy changes (shaded area is the 95% confidence interval). B) Phospho-occupancy time series of a multi-phosphorylated CaMKII-γ peptide. C) Reliable estimation of stable site occupancy is enabled by inducing dynamics with phosphatase treatment (see text, SI Appendix, Fig. S12). Examples of stoichiometry estimated with phosphatase treatment are shown here. Treated conditions are replicates of the 0 and 18 minute time points (boxed). D) Cumulative distributions of the initial phosphosite occupancies at 0 minutes (unactivated egg) classified by whether the sites increase, decrease, or are stable after egg activation. A uniform distribution would lie on the dotted line.

In total, we calculate confident stoichiometries (+/-25% occupancy of 95% confidence intervals) for ~500 sites (15% of dataset); ~150 of these are dynamic. With these data, we are able to compare stable versus regulated sites, which was difficult with previous approaches. For increasing, decreasing, or stable phospho-sites (classified by their relative trends), we show the cumulative distribution of their occupancy at 0 minutes (Fig. 5D). The majority of sites that increase have a low initial occupancy. The distribution of sites that decrease is skewed toward higher occupancy, though there is more density of lower occupied sites than reported in previous studies (24, 68, 69). Interestingly, the stable site distribution is the most highly occupied class. However, the difference between the distribution of stable sites and deceasing sites is perhaps more subtle than it appears. The apparent lack of intermediate occupancies is largely a product of applying a confidence interval cutoff to the data. The cutoff improves data quality, but disproportionally filters stable sites (SI Appendix, Fig. S13).

### Protein phosphorylation dynamics following egg activation

To explore the function of the regulated phospho-sites, we performed motif and gene set enrichment analysis (GSEA) on the dominant trends of the dataset: increasing and decreasing stoichiometries. While the trends we detect are a subset of the total sites in the egg, their signatures are nevertheless revealing of the classes of regulation that are occurring. We were careful to exclude misleading trends that occur during multisite phosphorylation where, for example, the increase of a single form is actually the loss of a double form (SI Appendix, Fig. S14, Methods). Since relative trends are sufficient for motif and GSEA, we used all the data to increase statistical power. The minimal motifs for Cdk1 and MAP kinases (S/T-P) explain ~70% of the decreasing sites (Fig. 6A). We show several examples of the absolute dynamics for these phospho-sites, calculated from our estimated site occupancies and protein concentrations (14). The majority of these show substantial loss of phosphate, which is consistent with the expected reversal of mitotic phosphorylation (Fig. 6B). The rates of the trends align with cell cycle events. The majority of proteins are dephosphorylated with a halftime of ~10 min (some with <5 min) with minimal delay after egg activation. This matches the timing of sister chromatid separation in anaphase (30, 70) and the half time of CyclinB1/2 degradation (Fig. 2A). GSEA of the dephosphorylated proteins are annotated with cell cycle, spindle and nuclear pore functions (Fig. 6A, specific examples in SI Appendix, Fig. S15). In parallel, we also see a strong signal for increasing phosphorylation. The (S/T-P) motifs are under-represented in this class. Instead, the minimal motif for the calcium-sensitive CaMKII is enriched, explaining 25% of the sites (Fig. 6C). Many of the increasing phosphorylation trends are fast, with half-times of 5 min or ~2 min (Fig. 6D). These are matched temporally with the calcium wave, cortical actin rearrangement (71–73), and exocytosis events (Fig. 2E). Indeed, the phosphorylated proteins are enriched for actin-binding GO terms; all of the members of this set are also annotated as “Cell Periphery” (Fig. 6C) (Table S5). Given the localization of these events to the cortex, their fast rates, and the calcium sensitivity of cortical granule exocytosis (9), the increasing phosphorylation may in part promote the release of protein into the media.

**Figure 6:**
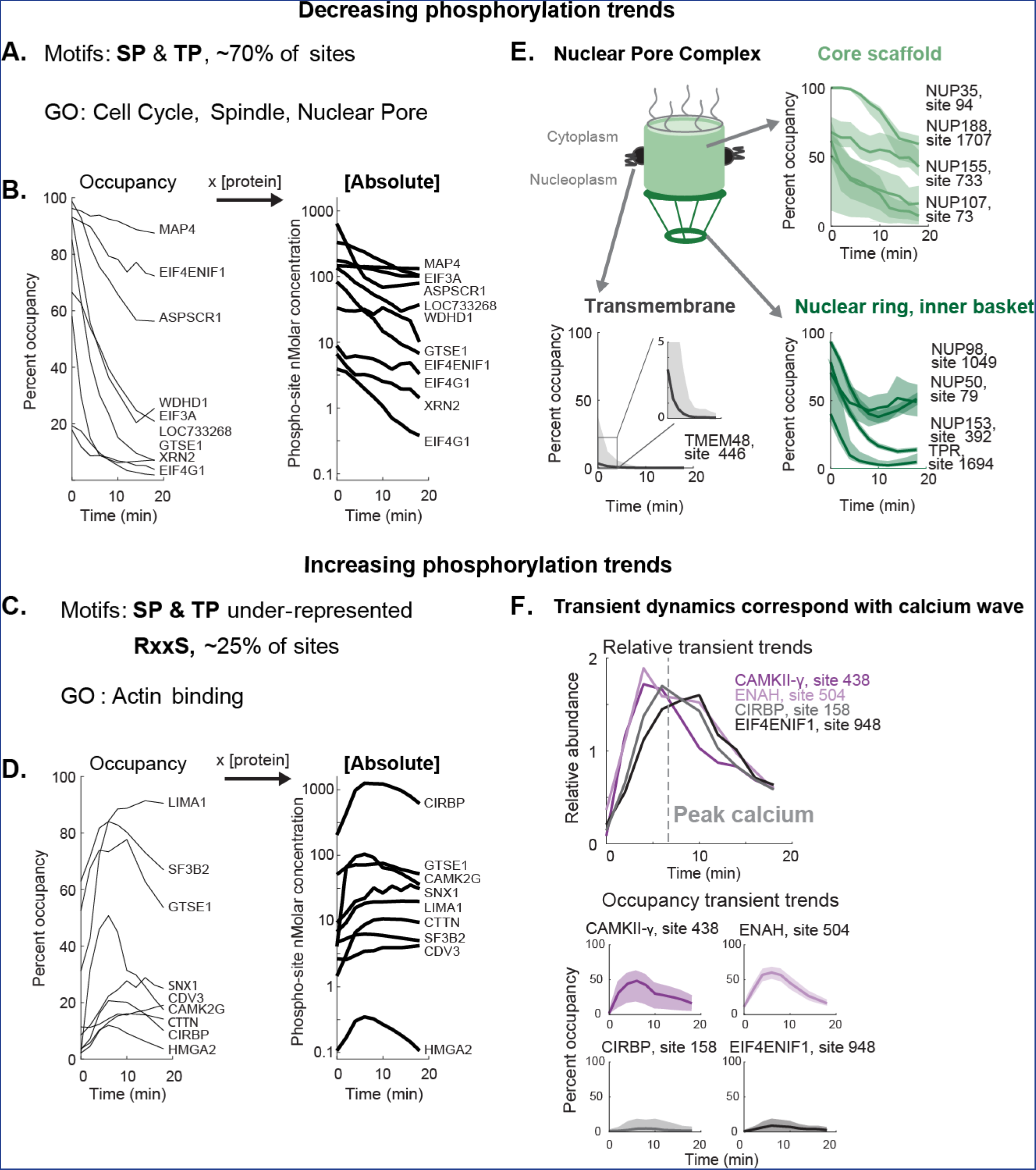
Phospho-site dynamics following egg activation. Analysis of the phosphorylation trends (increasing and decreasing) following egg activation with individual examples. A) Motif enrichment analysis (p <<0.01) results and gene set enrichment analysis (GSEA) of GO terms (p<0.01) results for dephosphorylated proteins. B) Time series of selected proline-directed phosphorylation plotted as occupancy and absolute phosphate dynamics (log10 transformed). All trends shown pass a 95% confidence interval width threshold of ±25%. C) Motif analysis results as in Fig. 6A for proteins with increasing phosphorylation trends and GSEA results. D) Examples of increasing phosphorylation (plotted as in Fig. 6B). E)Phospho occupancy trends showing differential dephosphorylation of Nuclear Pore Complex (NPC) regions with confidence intervals. F) Relative and phospho-occupancy time series of proteins showing transient trends corresponding to the peak of calcium concentrations.

### Nucleoporin dephosphorylation corresponds to order of NPC assembly

There are many intriguing vignettes in the data. One example is the differential rates of dephosphorylation of nuclear pore sub-complexes (Fig. 6E, SI Appendix, Fig. S16). During entry into mitosis, nucleoporin phosphorylation promotes Nuclear Pore Complex (NPC) disassembly (74). The sequence of post-mitotic NPC reassembly may be controlled by differential reversal of the mitotic phosphorylation (75). Our observations give some support for this hypothesis. Rates of dephosphorylation on nucleoporin phospho-sites cluster by regions (76) of the NPC (Fig. 6E) (discussed further in SI Appendix, Fig. S16). These rates mostly conform to the sequence of assembly (77, 78). The fast dephosphorylation (~2-5 min half times) of inner basket and nuclear ring components (e.g., NUP153) is consistent with their early roles in NPC assembly (79). The slower dephosphorylation (~10 min half time) of the Core Scaffold components (e.g., NUP188) is consistent with their later recruitment (SI Appendix, Fig. S16). One notable exception is that while the incorporation of NUP98 is reported to be an intermediate assembly step, it is dephosphorylated rapidly (4 min half time). Phosphorylation of NUP98 was previously shown to be rate limiting for NPC disassembly (80); it may be that its dephosphorylation is required or limiting for reassembly as well. There are two important caveats to the kinetic discrepancies: 1) nucleoporins or nucleoporin partial complexes may function outside the nuclear pore (e.g., NUP98 at the spindle) (81, 82); 2) the nuclear envelope in the egg is packaged in structures called annulate lamellae, which also repackage dynamically after fertilization (83). Our observations may reflect this repackaging, rather than the canonical post mitotic reassembly. For example, the large relative change but small stoichiometric decrease of the transmembrane nucleoporin TMEM48 may indicate a separate pool of molecules, perhaps on the annnulate lamellae (Fig. 6E). Nevertheless, these differential rates provide new information on how NPC reassembly may be regulated in the egg and by extension in other circumstances.

### Ca++ sensitive responses show substantial differences in occupancy

The data also offer insights into the pattern of calcium-initiated signaling. The calcium wave peaks in cytosolic concentration around 5 minutes post-fertilization and declines thereafter (84). Fig. 6F shows a set of relative phosphorylation changes from a larger cluster (Fig. 1E) that correlates with the calcium wave. A prominent example is CaMKII-γ, which is important for egg activation in the mouse (85), though the role of its phosphorylation is unclear. Three additional proteins with phospho-sites that correlate with the calcium wave are shown (Fig. 6F). While the relative changes are nearly overlapping, the stoichiometries are very different. The two translational regulators (CIRBP, EIF4ENIF1) show phospho-sites that change on a small fraction (<2%) of the proteins. In contrast, over half of the molecules of the gamma subunit of CaMKII and ENAH (an Ena/Vasp actin binding protein) are phosphorylated, supporting the hypothesis that the increasing phosphorylation class is related to calcium signaling and the remodeling of cortical actin. A possible explanation for the low stoichiometry of the changes on CIRBP and EIF4ENIF1 is that this site of modification plays no significant role in regulation; perhaps these sites are modified through promiscuous activity of calcium-dependent kinases (e.g., CaMKII, PKC). Alternatively, the modification may be highly localized.

### Absolute changes of protein phosphorylation, degradation, and release

The data allow comparison of the absolute changes in the diverse yet connected processes occurring in parallel at fertilization (Table 1). We estimate the total change resulting from protein degradation as ~300 nM (~0.01% of the total protein mass), which is dwarfed by a nearly 50-fold higher loss (~1%) from protein release (roughly consistent with older estimates (9, 86)). The total change in phosphorylation, which occurs on a diverse set of proteins from the cell cycle machinery to calcium-sensitive kinases, is ~7,000nM. This number is an underestimate, as we capture only a fraction of the total dynamics. Nevertheless, this is an order of magnitude higher than the changes due to protein degradation. Unlike protein degradation, where there was a strong correlation between low abundance and instability (Fig. 2D), we see large changes in phospho-occupancy for proteins that span a 1000-fold range of concentration (Fig. 6B, D). This is striking evidence that phosphorylation has the capacity to change the activity of many abundant proteins within minutes. Protein degradation, even with an active E3 ligase like the APC/C, can only work so quickly on a small number of low abundance proteins, which seem to be core to cell cycle progression.

**Table 1:** Measured absolute changes

| **Class of Dynamic** | **Absolute change, nM** | **Fraction of proteome, %** |
| --- | --- | --- |
| Secretion | 13,000 | ~1 |
| Protein Degradation | 300 | 0.01 |
| Phosphorylation | 6,800 | 0.3 |
| Decreased | 4,200 | - |
| increased | 2,600 | - |

## Discussion

We measured the absolute protein and phospho-site dynamics at fine time resolution following fertilization. The data reveal a small number of degraded substrates and substantial protein dephosphorylation related to the cell cycle. We also detect release of protein related to the slow block to polyspermy and parallel increases in phosphorylation, which may be linked to the calcium wave. To aid in interpreting the data, we developed a new method for estimating phospho-site stoichiometry using multiplexed MS.

The phosphorylation data are compatible with a recent cell cycle model, which describes a quantitative model for the cell cycle oscillator in the *Xenopus* egg (87). Earlier experiments demonstrated the oscillation (88, 89), but a satisfactory explanation of how it oscillated depended on several features, including a time delay (about 15min) between activation of Cdk1 and activation of APC/C and then a very small delay in the destruction of targets once APC/C is activated (<2min). Our measurements are strikingly consistent with fast Cdk1 inhibition, as we see immediate commencement of dephosphorylation at hundreds of proline-directed sites with no obvious delay. A discrepancy is that we see a small delay (~2-4 min) in the degradation of APC/C substrates like CyclinB1/2, which is presumably required to inactivate Cdk1. The rates of these early events may be underestimated if there is sufficient metachrony of the cell cycle in the large frog egg (45, 48), but the ordering of the observations discussed here should be preserved.

The substrate specificity of the APC/C is thought to be conferred by the cofactors Cdh1 and Cdc20. We show that all detected APC/C^Cdh1^ substrates are indeed stable in the frog egg, where Cdh1 is not expressed. The inhibition of APC/C^Cdh1^ promotes S-phase entry (90) in the conventional cell cycle; the absence of Cdh1 and resulting stability of its substrates may assist in the bypassing of the G1 phase to S phase during the cleavage divisions in the early embryo.

All detected degradation targets (e.g., Cyclin B1/2, EMI2) are present at low abundance, suggesting that the amount of protein degradation is small overall. However, if there is a global increase in protein synthesis at equilibrium with protein degradation, we would underestimate the flux of degradation. Protein synthesis at egg activation is reported in flies and mice (18, 91, 92), but our data shows no evidence for a burst of synthesis of any of the 8,700 proteins. Our previous work also showed that new proteins comprise a small fraction of the frog embryo even hours post fertilization (15). In principle, 2μM of new protein could accumulate in ~20 mins (~7μM/hr, a rate measured later in development) (93). If this occurred during the first cell cycle, we would expect to identify many of these proteins given our sensitivity limits (SI Appendix, Fig. S5).

In addition to measuring the extent of protein degradation and synthesis following fertilization, our proteome-wide approach unveiled unexpected components of the slow block to polyspermy. Early studies established that protease activity is essential for this event; the inhibition of proteases leads to polyspermy (94). Paradoxically, we found that eggs release multiple endogenous protease inhibitors in response to fertilization. Though protease inhibitors were recently found in the perivitelline space of *Xenopus* eggs (95), their function was not addressed. There are several possible functions for released inhibitors: 1) they could prevent activity of the proteases inside the cortical granules, 2) they might control the activity of the proteases post-release, 3) they may act to block the sperm acrosome reaction (96).

Our application of quantitative proteomics to fertilization highlighted several general considerations for interpreting MS data. We concluded that phosphorylation dynamics can confuse the analysis of protein trends, giving the appearance of change in stable proteins. Dynamics in any high stoichiometric modification (e.g., acetylation) could also cause this same ambiguity. Using the median of a protein’s peptides can mitigate but not will not eliminate specious trends caused by dynamic modifications. On the phospho-site level, a source of ambiguity is that a *decreasing* trend of a singly-phosphorylated form results from an *increasing* multi-phosphorylated form (SI Appendix, Fig. S14). This general possibility must be considered when interpreting the direction of a given phospho-site trend. This phenomenon was discussed previously (97); our data provides clear examples.

The utility of measuring phospho-occupancy is well-recognized (24, 25, 66). However, current approaches are not able to estimate occupancy for multi-phosphorylated peptides, stable sites, or sites measured with multiplexed proteomics. More fundamentally, these methods do not give statistical information (i.e., confidence intervals), which makes interpreting the estimates difficult. Our approach provides a unified analytical and experimental framework to address these limitations. Though we developed this approach to exploit the power of multiplexed proteomics, the advancements are compatible with other methodologies. For example, label-free phospho-proteomics typically have lower measurement precision but are conducive to high-throughput formats (98). The additional conditions, in principle, give increased statistical power with the use of the regression framework. The higher sensitivity of label-free and other techniques (99) could mitigate a limitation of this approach, which is the low efficiency of measuring both the unmodified and phosphorylated peptides. Other methods for calculating phospho-stoichiometry, such as motif-targeting proteomics (100), could also incorporate this framework. Finally, the principles of the calculation could be extended to acetylation or ubiquitination, though we expect more success if occupancies are often >1%. These approaches can increase the power of the proteomics for analysis of posttranslational modification in diverse settings.

## Materials and Methods

### Egg activation procedure

Female *X. laevis* were induced with 700 U HCG. After 14 hrs, eggs were harvested, washed with 1× MMR (Marc’s Modified Ringers), de-jellied with cysteine (2% w/v, pH 8.0), and then kept in dishes with agar beds to avoid the eggs sticking to the dish. Eggs were placed in open-faced gel box with a 3% agar bed in 0.1X MMR and electro-shocked by applying an electric field of ~3 Volts/cm for one second. Thirty eggs were collected every two minutes until 18 minutes. The excess media was removed and samples were flash frozen in liquid nitrogen. For the phosphatase treatment experiment, the 12 and 16 min time points were replaced with phosphatase-treated replicates of the 0 min and 18 min time points. Each biological replicate was performed on separate days with different frogs (termed EA, EAp2, EApR3, EAp5b, where EA stands for “Electro-Activation” and the ‘p’ denotes an experiment where phospho-enrichment occurred). The research with *X. laevis* was performed under the oversight of the Harvard Medical Area Institutional Animal Care and Use Committee.

### Egg media time series

Egg media was collected at the similar time points as Fig.1B following electro-activation, except with the 0 min time point taken in triplicate and 12 and 16 min time points skipped. The media was dried to concentrate the sample and processed for MS analysis. The experiment was performed as described above with the following exceptions: 1) Eggs were placed in 0.01X MMR, to reduce the salt concentrations after dry-down. 2) An additional ~10mM of calcium was added to the media to assist in any wound healing caused by the electroshock. 3) The electric field was doubled to ~6 Volts/cm to account for less conductive media, as eggs did not activate at 3 Volts/cm. 4) ~1,000 eggs were used instead of 300 to increase the signal of protein accumulating in the media. After electroshock, the media was kept well-mixed by gently rocking the gel-box on the long axis by hand (several methods were tested by mixing with dye). Each time point was taken by extracting 2 mL of media and flash freezing with liquid nitrogen. We replaced the volume of media removed for harvesting at each time point to keep overall volume constant throughout the experiment, and normalized afterward to account for the removal of mass in the media with each time point taken. We lyophilized the collected media samples to prevent unwanted proteolysis from the released proteases. The experiment was done in triplicate with three different clutches on the same day, and then processed in parallel as described in the MS sample preparation section below, except without the alkylation and protein precipitation steps.

### General sample preparation for mass-spectrometry

Samples were prepared essentially as previously described (14, 16). Thirty eggs per time point were snap frozen and lysed with 250 mM sucrose, 1% NP40 substitute (Sigma), 5mM EDTA (pH 7.2), 1 Roche complete mini tablet (EDTA-free), 20 mM HEPES (pH 7.2), 10 μM Combretastatin 4A, and 10 μM Cyochalasin D while frozen (14). All buffers were made with HPLC water. Eggs were lysed by pipetting up and down forty times with a 200 μL pipette tip, vortexed at maximum speed for 10 seconds, incubated on ice for 10 minutes, and again vortexed for 10 seconds. Lysates were clarified by centrifugation at 4,000 RCF at 4°C for 4 minutes in a tabletop centrifuge. The cytoplasmic and lipid layers were mixed by gentle flicking and removed from the pelleted yolk. To the lysate, HEPES (pH 7.2) was added to 100 mM, and SDS was added to 2% (w/v) to provide additional buffering capacity and to denature the sample. The samples were reduced with 5 mM DTT for 20 minutes at 60°C. Cysteines were alkylated with 15 mM NEM for 20 minutes at room temperature (RT). Excess NEM was reacted with an additional 5 mM DTT at RT. Proteins were isolated by methanol/chloroform precipitation (101). The protein pellet was resuspended (~5 mg/mL) in 6 M guanidine HCl and 50mM EPPS, pH 8.5 with gentle pipetting and heated to 60°C for five minutes. For approximately 100-400 μg protein per condition, the sample was diluted to 2 M guanidine with 5mM EPPS, pH 8.5 and digested with LysC (Wako Chemicals) at 20ng/μL at RT for 14 hours. Next, we diluted guanidine HCl to 0.5 M with 5mM EPPS, pH 8.5 and digested further additional LysC at 20 ng/μL for ~15 minutes at RT, then added 10 ng/μL of sequencing grade Trypsin (Roche) and co-incubated at 37°C for 8 hours in an incubator. For each sample ~100-400 μg of peptides were dried down in a SpeedVac and resuspended with 100-150 μL of 500 mM EPPS, pH 8.0 respectively. If sample was not at pH ~8.0, an additional ~25-50mM of HCl was added. For labeling, we added 15-50 μL of TMT stock solution (0.2 mg/40 μl ACN) to each sample and incubated at RT for 2 hours (10 μg:1.5 μL peptide to TMT). Thereafter, we quenched the reaction with 10 mM Hydroxylamine for 15 minutes. All conditions were combined, acidified by addition of phosphoric acid to 5%, and were clarified by spinning at 21K RCF for 20 minutes. Samples were subjected to C18 solid-phase extraction (50mg, SPE) (SepPak, Waters) to desalt and isolate peptides. To reduce sample complexity, peptides were resuspended in a 10 mM sodium carbonate buffer (pH 8.0), then fractionated by medium pH reverse-phase HPLC (Zorbax 300Extend-C18, 4.6 X 250 mm column, Agilant) using an acetonitrile gradient from 5% - 39%. With a flow rate of 0.8 mL/min, fractions were collected into a 96 well-plate every 38 seconds, and then pooled into 24 fractions by combining alternating wells from each column of the plate. Each fraction was dried and resuspended in 20 μL of 1% phosphoric acid. Peptides from each fraction were desalted and extracted once more with reverse-phase purification, resuspended in 10 μL 1% formic acid. ~4 μL per fraction were analyzed by LC-MS.

### Phosphatase treatment

To phosphatase treat some samples without affecting the untreated conditions after multiplexing or at any subsequent steps, we made use of a thermally unstable phosphatase, which can easily be inactivated. We used the temperature labile Shrimp Alkaline Phosphatase (Affymetrix, product #78390). To minimize volume added to the samples, the phosphatase was concentrated to ~4U / μL on a 5K Amicon Ultra filter, spun at 4°C. Since the supplied buffer contains Tris, which can interfere with TMT labeling, we exchanged this buffer with 10mM EPPS at pH 8.0 and stored the enzyme in 50% glycerol. We see no activity loss from the buffer exchange, as assayed by p-nitrophenyl phosphate (pNPP) hydrolysis. Following completion of trypsin digestion, we allowed the temperature to cool to RT then added 10mM MgCl 0.1mM EDTA, 5mM EPPS pH 8.5. To the two phosphatase treated conditions, we added enzyme at a ratio of 3:1 peptide to phosphatase units and incubated for 12 hours at RT without purifying the peptides away from the proteases. The incubation is done in the presence of proteases as the dephosphorylated peptide creates a better substrate and allows for proteolysis of an otherwise missed cleavage caused by the charge of the phospho-group. Allowing for further cleavage of dephosphorylated peptides mimics the effect of an endogenous phosphatase more closely; otherwise the forms with missed cleavages will have lower abundance in the untreated samples and confound analysis. We determined that at RT, LysC does not cleave the phosphatase, while Trypsin does but at a reduced rate. The 3:1 ratio used gives sufficient activity while avoiding peptides from the proteolysis of the phosphatase from dominating the MS signal. After incubation, samples were lyophilized to dryness, and resuspended per the TMT-MS protocol discussed above. Importantly, all samples were incubated for 5 minutes at 65° C to destroy phosphatase activity in a water bath (inhibition under these conditions was confirmed beforehand with pNPP activity assays). Samples were cooled to RT before proceeding with the MS sample preparation.

### Phospho-peptide enrichment

We chose to multiplex peptides before the enrichment column to improve the data quality. There is a tradeoff of decreased yield and therefore depth, as it is not economical to label more than a few milligrams of material. We used 2.5-4 mgs of TMT-labeled peptides per replicate, enriched with 5 μm Titanium Dioxide microspheres (GL Sciences 5020-75000) and fractioned as previously described (102). A typical yield of 50-80 μg of peptides eluted from the column, with a median specificity of ~80%. Additional experimental details are provided in the SI Appendix.

### LC-MS analysis

Analysis performed essentially as previously described, with the spectra mapped to the PHROG reference database (14, 16). The main modification was the use of 5-notch MS3 on the protein-level and 3-notch MS3 on the phospho-level (103). Phosphorylation searches were performed with differential modification of +79.9663304104 on Serine and Threonine. Tyrosine was excluded as we found the majority of the identified tyrosine sites were erroneous identifications. Phospho-sites were localized with in-house software based on Ascore (104). Peptides with multiple phospho-sites are reported as “composite sites” and noted with a “;” delimiting each modified residue. All searches (for the protein replicates, phospho replicates, media, ubiquitin pull downs) were mapped to a single Protein Map and subjected to one Linear Discriminator/Protein Sieve Analysis to control the false discovery rate.

### Estimating a false discovery rate for protein loss

The full methodology is detailed and visualized in SI Appendix Methods and Fig. S2. The false discovery rate cutoff was calculated by comparing the geometric distance of the experimental data along with randomized data to an idealized degradation trend. The FDR is defined as the chance that a random trend could appear at a certain cosine distance from an idealized degradation trend.

### Classification of released proteins and novel APC/C substrates

We classified a protein as released (rather than degraded) either by direct experimental evidence, annotation, or evidence in the literature. In most cases, a protein that decreased in the egg could be clearly detected as increasing in abundance in the media. There were several cases where a protein decreased in the egg that we did not detect in the media, but these were clearly annotated as secreted proteins or were of a similar class as proteins for which we had direct evidence. For example, we infer the Exocyst gene family are released into the media because vesicle trafficking and components of the exocytosis machinery were detected recently in the perivitelline space in frogs (95). To be classified as a novel APC substrate, we imposed the following criteria: the protein passes a 1% FDR threshold for decreasing abundance, had no evidence (direct or from literature) of release from the egg, no evidence of a specious trend from phosphorylation (this was established by looking for reciprocal phosphorylations, or more generally, whether all the detected peptides behaved similarly), and was ubiquitinated.

### Phospho-stoichiometry regression-based algorithm

The full principle and methodology is included in the SI Appendix. The MATLAB implementation is freely available at https://github.com/marcpresler/OccupancyMS. The Total Least Squares regression package (105) is adapted from File ID: #31109 on MathWorks File Exchange.

### SI table guide

All tables are provided as separate sheets in a single excel file. Table S1 shows the rank-ordered degradation candidates as unique protein IDs with protein sequence. Table S2 shows the absolute loss of candidates passing a 1% FDR cutoff averaged by unique gene symbol. Table S3 classifies Table S2 by their mechanism of loss. Table S4 shows the proteins that apparently increase in abundance. Table S5 contains the Go Term enrichment output. Table S6 provides the raw data output for all experiments. Table S7 and S8 give the normalized protein and phospho-site data with replicates averaged. Table S9 contains the phospho-site occupancy data.

### Methods in SI Appendix

EM imaging time series, data normalization, calculating absolute changes, estimating maximal degradation rates of the APC/Proteasome, K-means clustering, multi-site artifact correction, motif and gene set enrichment analysis.

## Author contributions

Conceptualization: MP, MW, MWK. Experiments: MP, EV, MW, PC, RK. Analysis: MP, EV, MW, LP. Stoichiometry method: MP, AMK, EV, MW. Writing: MP, EV, MW, AMK, MWK. Supervision: MW, SPG, MWK.

## Acknowledgements

This work was supported by NIH grants HD091846, HD073104, GM103785 to MWK, GM39565 to TJM, and Burroughs Wellcome Fund and Mallinckrodt Award to AMK. We thank Chris Rose and Joao Paulo for help with mass spectrometers and the Gygi lab computational team for bioinformatics support, Ying Lu, Tao Wu, Tim Mitchison, and Angela DePace for helpful discussion, Katy Hartman, John Ingraham, and Randy King for insight and inspiration toward the stoichiometry calculation, and Becky Ward for critical comments on the manuscript. We thank the PRIDE team for proteomic data distribution.

## Accession Numbers

The mass spectrometry proteomics data have been deposited to the ProteomeXchange Consortium via the PRIDE (106) partner repository with the dataset identifier PXD006639.

